# The crystal structure of a Polerovirus exoribonuclease-resistant RNA shows how diverse sequences are integrated into a conserved fold

**DOI:** 10.1101/2020.04.30.070631

**Authors:** Anna-Lena Steckelberg, Quentin Vicens, David A. Costantino, Jay C. Nix, Jeffrey S. Kieft

## Abstract

Exonuclease-resistant RNAs (xrRNAs) are discrete elements that block the progression of 5’ to 3’ exonucleases using specifically folded RNA structures. A recently discovered class of xrRNA is widespread in several genera of plant-infecting viruses, within both noncoding and protein-coding subgenomic RNAs. The structure of one such xrRNA from a dianthovirus revealed three-dimensional details of the resistant fold but did not answer all questions regarding the conservation and diversity of this xrRNA class. Here, we present the crystal structure of a representative polerovirus xrRNA that contains sequence elements that diverge from the previously solved structure. This new structure rationalizes previously unexplained sequence conservation patterns and shows interactions not present in the first structure. Together, the structures of these xrRNAs from dianthovirus and polerovirus genera support the idea that these plant virus xrRNAs fold through a defined pathway that includes a programmed intermediate conformation. This work deepens our knowledge of the structure-function relationship of xrRNAs and shows how evolution can craft similar RNA folds from divergent sequences.

## INTRODUCTION

RNA structure-dependent exoribonuclease resistance is now well-established as a means of subgenomic RNA production or maintenance by many viruses (Slonchak and Khromykh 2018). This mechanism depends on discrete, specifically folded RNA elements in viral genomes called xrRNAs (for ‘e<u>x</u>oribonuclease-resistant <u>RNAs</u>). xrRNAs block the 5’ to 3’ progression of processive cellular exoribonucleases using only a folded RNA element, without accessory proteins (Chapman et al. 2014a). In so doing they protect downstream RNA from degradation. xrRNAs were discovered in flaviviruses and have since been identified in other viruses, including plant viruses of the *Luteoviridae* and *Tombusviridae* families (Pijlman et al. 2008; Steckelberg et al. 2018a; Iwakawa et al. 2008). In flaviviruses, xrRNAs have been found exclusively at the beginning of the 3’ untranslated region (UTR) of the viral genome where they function in the generation of viral noncoding RNAs, whereas in *Luteoviridae* and *Tombusviridae*, xrRNAs are associated with both noncoding and protein-coding regions of the viral genome (Fig. 1A) (Steckelberg and Vicens et al. 2018b). Despite evidence that xrRNAs may be widespread and involved in a variety of viral processes, we are only beginning to understand the principles of xrRNA folding that lead to structures capable of blocking host cell exoribonucleases.

**Figure 1:**
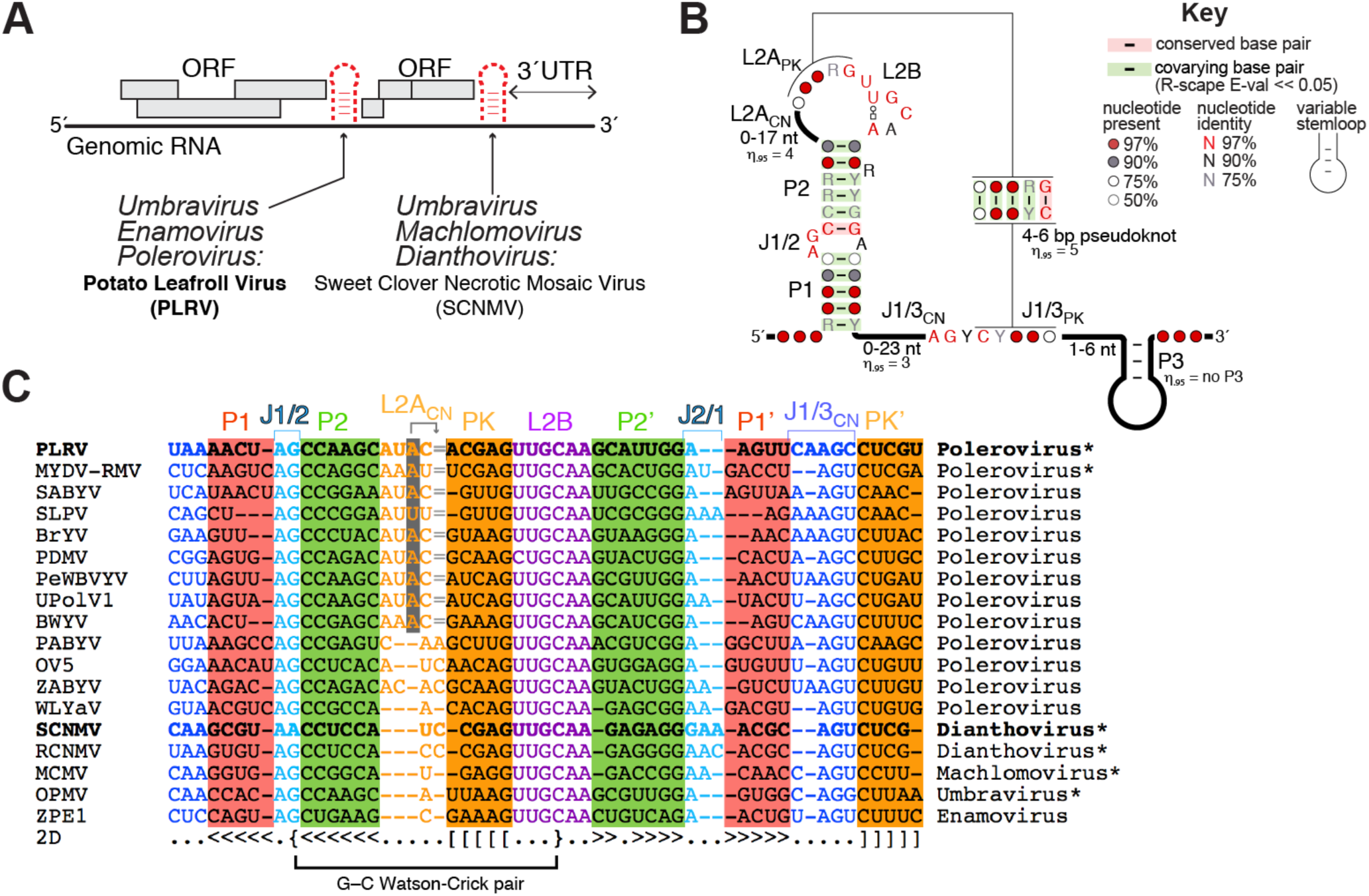
A brief overview of plant xrRNAs. (A) Depending on the plant virus family, xrRNAs are located within intergenic regions and/or at the beginning of the 3’ UTR. (B) Consensus secondary structure for xrRNAs_LT_. The 95^th^ percentile is indicated for all variable regions. Covariation analysis was expanded and updated from (Steckelberg and Vicens et al. 2018b). (C) Alignment of xrRNA_LT_ sequences from select plant viruses. Bold, sequences for which a crystal structure is available; *, xrRNA confirmed to be resistant to Xrn1 *in vitro*; grey highlight in L2A_CN_, the highlighted nucleotide aligns two nucleotides downstream in three dimensions, as indicated by the arrow pointing to the ‘=’ sign. The G–C pair involving J1/2 and L2B is labeled. The complete alignment is shown in Supplemental Fig. S1.

Crystal structures of xrRNAs from flaviviruses revealed that two pseudoknots and other long-range interactions create a ring-like fold that wraps around the 5’ end of the xrRNA (Chapman et al. 2014a; Akiyama et al. 2016). Mechanistic studies suggest that this ring protects downstream RNA by ‘bracing’ against the surface of the exoribonuclease, acting as a physical block to the 5’ to 3’ progression of the enzyme (MacFadden and O’Donoghue et al. 2018; Chapman et al. 2014a; Akiyama et al. 2016). A recent crystal structure of an xrRNA from Sweet Clover Necrotic Mosaic virus (SCNMV; a dianthovirus from the *Tombusviridae* family) showed a different three-dimensional fold conferring exoribonuclease resistance (Steckelberg et al. 2018a). Specifically, the SCNMV xrRNA was captured in a folding intermediate state which contains a Watson-Crick base paired helix (P1) that must be unwound to allow the formation of a pseudoknotted ring structure. Single-molecule Förster resonance energy transfer (smFRET) experiments revealed that the SCNMV xrRNA’s intermediate state can be remodeled by the helicase activity of the arriving exoribonuclease to favor the resistant pseudoknotted fold (Steckelberg et al. 2018a). While a ring encircling the 5’ end seems to be thus far a defining characteristic of all xrRNAs, the degradation-induced structural remodeling seen in SCNMV’s xrRNA appears to be unique. Overall, the xrRNAs of the flavivirus-type (xrRNA_F_) and the dianthovirus*-*type (xrRNA_D_) are distinguishable based on their different underlying three-dimensional folds.

The discovery of intramolecular interactions within xrRNA_D_ that could not be predicted from the sequence informed subsequent bioinformatic-based approaches that identified >40 new examples of this class of xrRNA (Steckelberg and Vicens et al. 2018b). These xrRNAs pervade the *Luteoviridae* and *Tombusviridae* families in general, and the *Umbravirus* and *Polerovirus* genera in particular (polerovirus sequences make for ∼ 2/3 of this alignment). Some of the deadliest viruses in agriculture contain an xrRNA, such as Potato Leafroll Virus (PLRV; leading responsible virus for worldwide potato yield loss (Wale et al. 2008)), Maize Chlorotic Mottle Virus (MCMV; responsible for 90% maize/corn yield loss in sub-Saharan Africa (Mahuku et al. 2015)) and Maize Yellow Dwarf Virus-RMV (MYDV-RMV, formerly BYDV-RMV (Krueger et al. 2013), responsible for 30% yield loss in affected winter wheat fields (Perry et al. 2000)). All newly identified xrRNAs in *Luteoviridae* are found in intergenic regions, whereas xrRNAs from *Tombusviridae* are more generally found at the beginning of the 3’ UTR (Steckelberg and Vicens et al. 2018b). Because they are widespread in the *Luteoviridae* and *Tombusviridae*, we refer to this class as xrRNA_LT_ in this manuscript rather than xrRNA_D_. Interestingly, the many newly-discovered xrRNA_LT_s contain conserved sequence elements absent from the crystallized SCNMV xrRNA (Fig. 1B,C; (Steckelberg and Vicens et al. 2018b)). This raised the questions of whether the newly identified xrRNAs_LT_ form the same fold as the dianthovirus xrRNA and if so, how they accommodate these different sequence elements.

To address the universality of the interactions necessary to support an xrRNA_LT_ fold as first seen in SCNMV, we explored the structure and function of the xrRNA_LT_ from PLRV using x-ray crystallography and *in vitro* assays of exoribonuclease resistance. In the crystal, the PLRV xrRNA_LT_ adopts a folding intermediate conformation similar to that of the SCNMV xrRNA_LT_. But as the PLRV xrRNA_LT_ is ∼ 10% longer than the SCNMV xrRNA_LT_, our structure reveals how extra nucleotides leading to extended helices and loops generate interactions and motifs not seen in the SCNMV structure. Using site-directed mutagenesis coupled with functional assays, we show how these interactions are critical for exoribonuclease resistance. Together, our results rationalize sequence conservation patterns of xrRNA_LT_, thereby providing insights into the mechanism of programmed exoribonuclease resistance in the viral world.

## RESULTS AND DISCUSSION

### Crystallization of a polerovirus xrRNA_LT_

The previously-reported structure of the SCNMV xrRNA_LT_ revealed how an RNA sequence of only 44 nucleotides adopts a specific fold to generate an exoribonuclease-resistant structure (Steckelberg et al. 2018a). Because the dianthovirus sequences in the original xrRNA_LT_ alignment were not representative of the broader diversity in the expanded alignment (Fig. 1C, Supplemental Fig. S1), the SCNMV structure could not fully explain the consensus secondary structure (Fig. 1B). In particular, all xrRNAs_LT_ contain a guanosine in the J1/2 internal loop between the P1 and P2 stems (G8 in PLRV; previously referred to as L1; Fig. 1C,2A), except for SCNMV which contains an adenosine. In SCNMV, this A is involved in long-range interactions with the highly conserved L2B region that could not be supported by a G. Additionally, xrRNA_LT_ sequences are longer for poleroviruses than for dianthoviruses, with extra nucleotides in P2 and in the less-conserved regions within the apical loop L2A (including the part involved in the pseudoknot) and the joining region J1/3 (Fig. 1B,C). Examining the SCNMV structure did not suggest how these extra nucleotides could be accommodated. Finally, the SCNMV xrRNA_LT_ was crystallized in a state that appeared to be a necessary folding intermediate. Additional structural information was needed to confirm this as an authentic intermediate or to visualize the final pseudoknotted state.

**Figure 2:**
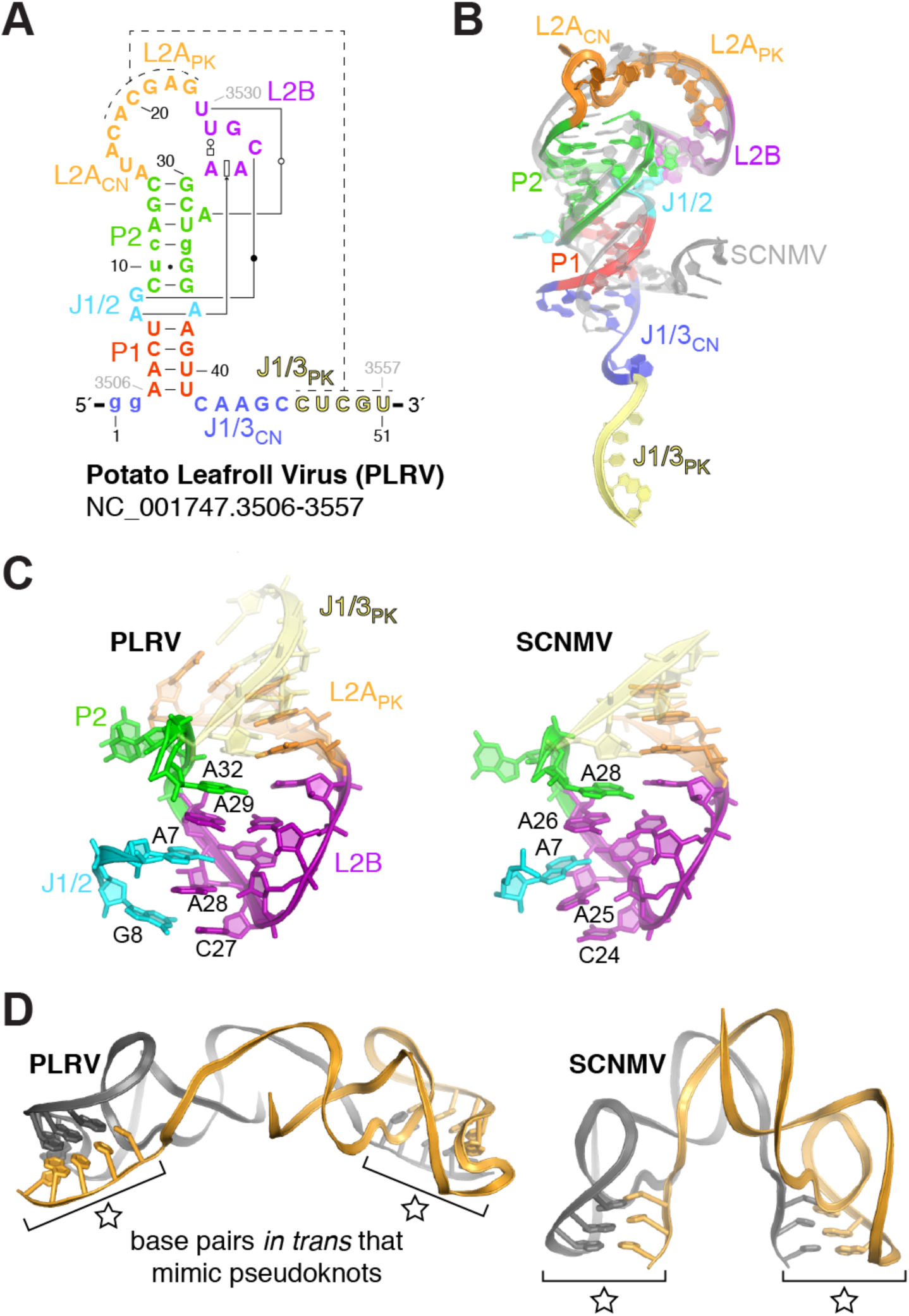
Structure of the PLRV xrRNA. (A) Secondary structure of the crystallized PLRV RNA, with matching colors to Fig. 1C. Engineered nucleotides (see Methods) are shown in lower case. Non– Watson–Crick base pairs are according to the Leontis–Westhof annotation (Leontis and Westhof 2001). (B) Ribbon representation of the PLRV xrRNA_LT_ structure in colors to match (A), over that from SCNMV in grey. (C) Side-by-side comparison of the conserved structural core involving J1/2, L2B and P2. (D) The PLRV and SCNMV xrRNA_LT_s both crystallize as a domain-swap dimer, in which the intramolecular pseudoknot is replaced by intermolecular base pairing (indicated by a star).

To allow a structural comparison with the SCNMV xrRNA_LT_, we selected several diverse xrRNAs_LT_ that had the conserved G, expanded regions in P2, L2A and J1/3, and various pseudoknot lengths (4–5 base pairs). Of these, the xrRNA_LT_ from PLRV yielded crystals that diffracted to 5–7 Å on our home source but that we could not reproduce (construct #1, Supplemental Fig. S2A). Diffraction was improved to ∼3 Å upon adding of 100 mM guanidinium hydrochloride during crystallization (see Methods), and introducing a G.U wobble base pair ‘phasing module’ at various locations to aid with phasing (Keel et al. 2007) (Supplemental Fig. S2A). Introducing the phasing module in constructs #1–4 did not alter the ability of the RNA to resist exonuclease-catalyzed degradation (Supplemental Fig. S2B). The structure of construct #3 was eventually solved to 2.6 Å from combined datasets collected at a synchrotron, using single anomalous dispersion phasing at the L-I edge of iridium (Supplemental Table S1; Supplemental Fig. S3) (Batey and Kieft 2016).

### A common intermediate fold for xrRNA_LT_ elements

The global structure of the PLRV xrRNA_LT_ is similar in shape and architecture to that of the SCNMV xrRNA_LT_. Like SCNMV, the PLRV xrRNA_LT_ folds as a “bent-over” stem-loop in which L2B creates a docking point for long-range tertiary interactions involving nucleotides located in J1/2 (Fig. 2A,B). The all-atom root-mean square deviation (RMSD) is 0.9 Å for a superimposition of L2A_PK_ (the section of L2A involved in <u>p</u>seudo<u>k</u>not formation; to be distinguished from L2A_CN_, the 5’ part of L2A <u>c</u>o<u>n</u>necting P2 to L2A_PK_) and L2B from SCNMV and PLRV, indicating that the two RNAs share a similar “core” structure (Fig. 2C). The improvement in diffraction quality upon addition of guanidinium is likely due to binding around J1/2 and J2/1, at sites where iridium(III) hexamine complexes bind in the SCNMV structure (Supplemental Fig. S3C). The guanidinium-RNA interaction will be described in more depth elsewhere.

As in the SCNMV xrRNA_LT_, the PLRV xrRNA_LT_ was crystallized in a ‘domain-swap dimer’ conformation, in which the intramolecular pseudoknot required for activity was replaced by interactions *in trans* between the two molecules in the asymmetric unit. Five Watson-Crick base pairs formed between the L2A_PK_ region of one molecule and J1/3_PK_ of the second molecule (Fig. 2C,D). Hence, the PLRV RNA was captured in a similar folded state to the SCNMV RNA (Steckelberg et al. 2018a), although the relative angle between the two molecules is about three times wider for PLRV (Fig. 2D). Because the P1 stem formed, but not the pseudoknot, in both SCNMV and PLRV xrRNA_LT_, this “open” state is likely an authentic intermediate conformation adopted by diverse xrRNAs_LT_ to ensure that the correct topology is ultimately achieved (Steckelberg et al. 2018a). This conclusion is further supported by previous smFRET and functional studies which showed that mutations favoring this intermediate state are not deleterious to exonuclease resistance, while those destabilizing this state reduce exonuclease resistance (Steckelberg et al. 2018a). Taking into account the sequence and secondary structure conservation of known xrRNAs_LT_ (Fig. 1B,C; Supplemental Fig. S1), we can expect that every xrRNA_LT_ generally follows a similar folding strategy, in which the initial formation of P1 positions the 3’ end relative to the 5’ end of the RNA, and thereby ensures that adoption of the final “closed” pseudoknotted form creates a protective ring encircling the 5’ end. As previously proposed, this closed form, not yet directly visualized, would be promoted by the helicase activity of the approaching exoribonuclease, which unwinds P1 (Steckelberg et al. 2018a).

### An unexpected G–C base-pair contributes to the ‘bent over’ conformation

The PLRV and SCNMV xrRNA architectures are globally similar (Supplemental Fig. S4), however several local structural differences account for the diversity in xrRNA_LT_ sequences. Although A8 is paired with G34 in SCNMV, the > 97% conserved G8 and A37 at the equivalent positions in PLRV and all other xrRNAs_LT_ identified are both bulged out from the stem (Fig. 3A). A37 forms crystal contacts (Supplemental Fig. S3D), but G8 pairs with C27 in L2B (equivalent to C24 in SCNMV; Fig. 2C,3A). The G8-C27 pair caps a stack of purines from L2B, J1/2 and P2 that is otherwise conserved between SCNMV and PLRV (Fig. 2C). Because the overall conformation of L2B relative to P1 and P2 is similar between the two structures, this additional base pair could be viewed as stabilizing but not altering the xrRNA_LT_ conformation (Fig. 2C,3A). The local changes at J1/2 and L2B seen in the PLRV structure may seem minimal, but they contribute to explaining the > 97% sequence conservation at J1/2 and L2B in a manner that the SCNMV structure could not.

**Figure 3:**
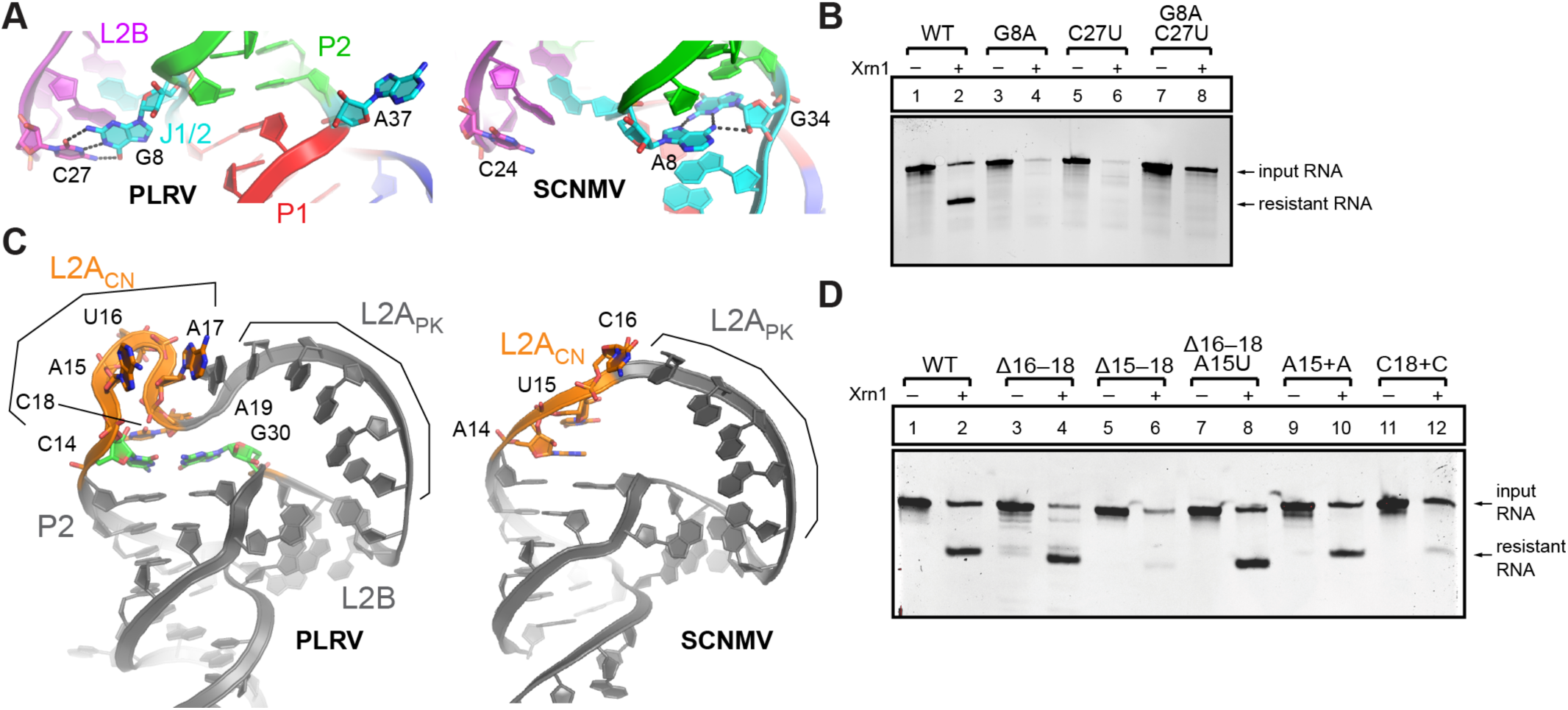
Structural diversity of the xrRNA_LT_ fold. (A) Sequence differences at J1/2 lead to the formation of the G8-C27 pair with L2B in PLRV (left; color coding as in Fig. 1C). Similar view for SCNMV (right). (B) *In vitro* Xrn1 degradation assay of PLRV xrRNA_LT_ WT and G8-C27 mutants. (C) Architecture of the L2A_CN_ and L2A_PK_ regions in PLRV (left) and SCNMV (right). (D) *In vitro* Xrn1 degradation assay of PLRV xrRNA WT and L2A_CN_ mutants.

The high conservation of the G8-C27 pair suggests it is important for exoribonuclease resistance. To test this, we generated two mutant RNAs that each disrupted the ability of these two bases to form a Watson-Crick base pair: G8A and C27U (Supplemental Fig. S5). When challenged with the recombinant 5’ to 3’ exoribonuclease Xrn1 in our established exoribonuclease resistance assay (Chapman et al. 2014b), both mutant RNAs were degraded, indicating they are unable to block progression of the nuclease (Fig. 3B, lanes 3-6). A double mutant G8A+C27U was designed to restore the ability of the Watson-Crick pair to form (Supplemental Fig. S5). Yet, this mutant was also unable to block progression of the exoribonuclease (Fig. 3B, lanes 7-8), suggesting that the specific identity and configuration of the base pair is important. This result is consistent with the sequence alignment, which shows no covariation at these positions (Supplemental Fig. S1). Substitution at these positions likely results in local structural changes that prevent formation of this tertiary base pair. Hence, the conservation of G8 and C27 serves not only to create a base pair, but also to promote structures that make these two bases available to pair. The fact that this pair is not present in SCNMV suggests it may not be required in all contexts, as evolution can craft compensatory stabilizing interactions.

### PLRV’s longer sequence is accommodated within the active fold

The P2 stem and the predicted pseudoknot are longer by one base pair in PLRV compared to SCNMV. L2A_CN_ is also longer (by one nucleotide) in PLRV compared to SCNMV. Finally, the J1/3 connector (J1/3_CN_) is longer by two nucleotides (Fig. 1C; next section). The PLRV xrRNA structure shows how these additional nucleotides are accommodated. In the SCNMV structure, L2A_CN_ has a 5’-AUC-3’ sequence, while PLRV has 5’-AUAC-3’, which leads to a significant structural adjustment. For SCNMV, A14 and U15 stack against P2 and C16 stacks against the pseudoknot-like structure formed *in trans*. For PLRV, the equivalent A15 and U16 flip out in opposite directions, and C18, equivalent to C16 from SCNMV, stacks against the elongated P2 (Fig. 3C, Supplemental Fig. S3D). Spatially, C14 from the C-G pair extending P2 and C18 in PLRV are in similar locations to A14 and U15 in SCNMV. The protrusion comprising A15-A17 in PLRV is accommodated by stacking of A15 and A17 against L2A_PK_ (via A19; Fig. 3C) and A37 from a symmetry-related molecule (Supplemental Fig. S3D). U16 is in the vicinity of another symmetry-related molecule, although the poor quality of the map for that residue suggests it is dynamic (Supplemental Fig. S3D). This side-by-side comparison of the xrRNA_LT_ structures from SCNMV and PLRV reveals extensive differences that were not apparent from sequence alignments but that show how differences in the sequences are accommodated in a globally similar fold.

To test whether the alternative stacking strategy against L2A_PK_ involving a distorted backbone of L2A_CN_ is required for resistance, we tested xrRNA mutants that alter L2A_CN_ (Supplemental Fig. S5). Removal of nucleotides 16–18 in PLRV to return to an SCNMV-like L2A_CN_ does not affect resistance, while additionally removing A15 does (Fig. 3D, lanes 3–6). Most likely, removal of A15 would make L2A_CN_ too short to support the proper conformation of L2A_PK_ and L2B. Replacing A15 with a U in the 16– 18 deletion mutant does not affect resistance (Fig. 3D, lanes 7–8), suggesting that the presence of a nucleotide at that position is more important than its identity. Notably, xrRNA_LT_ tolerates the presence of an extra nucleotide in L2A_PK_ after A15, but not after C18 (Fig. 3D, lanes 9–12). A straightforward interpretation of these results is that stacking of A17 and C18 is important to support the active fold, while stacking of A15 may only occur due to the presence of A37 from a symmetry-related molecule. Overall, this analysis helps to make sense of the generally longer L2A_CN_ and P2 stems in poleroviruses compared to dianthoviruses (Supplemental Fig. S1).

### Conserved interactions in the domain-swap dimer have a functional role

PLRV and SCNMV xrRNA_LT_ both crystallized in open conformations in which the pseudoknot is replaced by similar base pairs *in trans* (‘domain-swapped dimer state’; Fig. 2D). The intermolecular dimer contacts likely created favorable crystal packing, and dimerization is favored at high RNA concentrations used for crystallization. Using electron mobility shift assays (EMSA) we showed that at the RNA concentration of 150 μM used for crystallization, over 60% of the RNA molecules form dimers. At a lower RNA concentration used for functional *in vitro* assays (∼6 μM), RNA molecules are predominantly monomeric, suggesting that dimerization is a crystallization artifact and not the functional state of the RNA (Supplemental Fig. S6). Supporting this conclusion, the relative orientation of the two RNA molecules in a dimer is different in PLRV and SCNMV xrRNA crystals (Fig. 2D). The intermolecular base pairs involving L2A_PK_ of one molecule and J1/3_PK_ of the other are nonetheless similarly mimicking the pseudoknot in both structures (Fig. 2C,D).

The interactions involving J1/3_CN_ (42–46 in PLRV; Fig. 2A) immediately adjacent to J1/3_PK_ are also conserved in the PLRV and SCNMV structures (Fig. 4A–D). Examination of our sequence alignment reveals that these nucleotides are highly conserved, with > 97% conservation of A44 and G45 (A40 and G41 in SCNMV), and the conserved presence of a pyrimidine at position 46 (Fig. 1C). A44, G45 and U46 all stack together and help bridge the bent-over appearance of this hairpin. In particular, A44 interacts with the Hoogsteen side of A7 and is part of an interaction network involving U25 and G26 from L2B. G45 stacks on A44 and bridges A7 to the conserved A29-U25 pair (Fig. 4C). Finally, C46/U42 points toward a negatively charged region formed by the backbone of P1 and P2, where iridium(III) hexammine complexes have been assigned in both PLRV and SCNMV (Fig. 4D). The absence of a purine at this position across xrRNA_LT_ sequences (Supplemental Fig. S1) could be justified by the steric clashes or the disturbance of potential physiological metal ion binding sites that a purine would cause.

**Figure 4:**
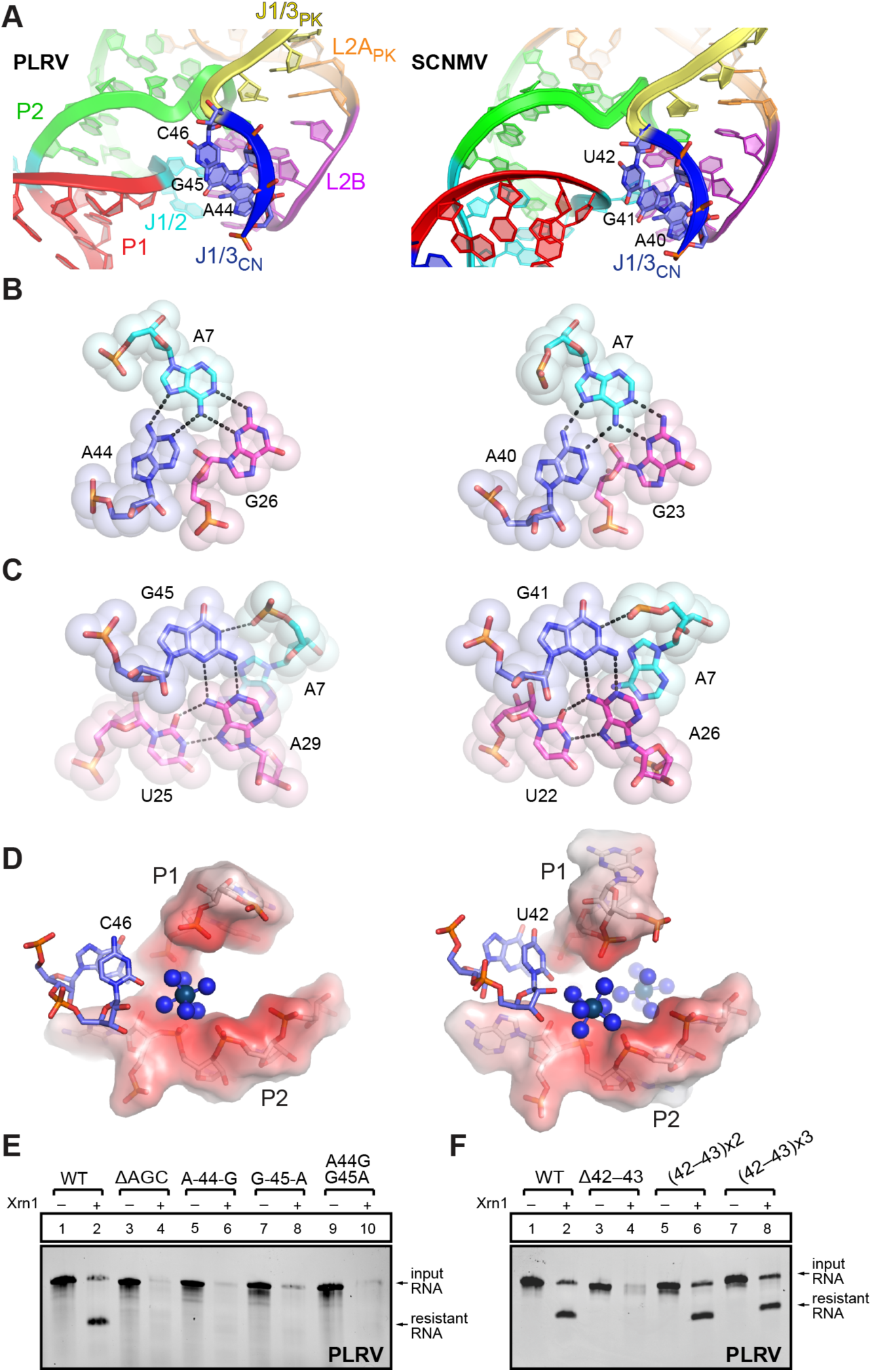
Conserved interactions involving J1/3_CN_ at the dimer interface. (A) Close-up on the region involving J1/3_CN_ (A44, G45, C46 in PLRV, and A40, G41, U42 in SCNMV) in both structures. For panels A-D, the PLRV structure on the left side is compared to the SCNMV on the right side. (B) Interaction network involving A44. (C) Interaction network involving G45. (D) C46/U42 are in the vicinity of a negatively charged region (in red) formed by the backbone atoms of P1 and P2. (E–F) *In vitro* Xrn1 degradation assays of PLRV xrRNA WT and J1/3_CN_ mutants.

To interrogate the importance of the conservation at J1/3_CN_, we generated mutants to disrupt the interaction involving nucleotides 44–46 (Supplemental Fig. S5) and we tested them for exoribonuclease resistance. As expected, deletion of A44–C46 (ΔAGC) removes the ability to resist degradation by Xrn1 (Fig. 4E, lanes 3–4). But even purine to purine substitutions at positions 44 and/or 45 are sufficient to disrupt tertiary interactions, leading to complete degradation by Xrn1 (Fig. 4E, lanes 5–10). In conclusion, the conserved 5’-AGY-3’ sequence in J1/3_CN_ is involved in tertiary interactions that are key for resistance against Xrn1 degradation. This suggests that the interactions occurring *in trans* within the dimer probably occur *in cis* to stabilize the active fold.

The equivalent of J1/3_CN_ in flavivirus xrRNA is part of the ring-like structure that wraps around the 5’ end (Steckelberg et al. 2018a; Chapman et al. 2014a), which suggests its length would be important for proper ring formation and hence resistance. Because J1/3_CN_ shows variation in length across xrRNA_LT_ elements beyond the more frequent length of 6 nucleotides (Supplemental Fig. S1), we aimed to test the effect on resistance of decreasing or increasing J1/3_CN_. As for P2 and L2A_CN_ discussed above, J1/3_CN_ is on the shorter side for SCNMV (3 nucleotides vs. 5 in PLRV; Fig. 1C). Shortening the J1/3_CN_ in the PLRV xrRNA_LT_ as with the ΔAGC mutant affected function (Fig. 4E), although that effect is more likely due to the disruption of the associated interaction networks (Fig. 4A–D). Upon shortening J1/3_CN_ by two nucleotides upstream of the AGC stretch (deleting C42 and A43), resistance to Xrn1 was lost (Fig. 4F, lanes 3–4). This result suggested that this mutant may no longer be able to form a ring that would prevent it from being degraded. In contrast, extending J1/3_CN_ by 2 nucleotides [(42-43)x2] or 4 nucleotides [(42-43)x3] did not significantly reduce exoribonuclease resistance (Fig. 4F, lanes 5–8; Supplemental Fig. S5), indicating a tolerance for a somewhat longer J1/3_CN_, in line with the observed variation across xrRNA_LT_ sequences.

### Outlook

The presence of xrRNAs in a growing list of viruses from several major superfamilies, combined with emerging structural information, continues to expand our insight into their diversity and evolution. The work presented here further shows that all xrRNA structures solved thus far have the potential to form a ring-like fold, but there are different ways to achieve it. In that sense, visualizing the closed conformation for an xrRNA_LT_ —i.e., with a ring encircling the 5’ end— remains a priority as it would enable a thorough comparison across the currently known xrRNA active folds. The fact that xrRNAs from flaviviruses and dianthoviruses/poleroviruses are variations on the same theme suggests they may have evolved from a common precursor. Indeed, an emerging hypothesis in biology is that all viruses have a common ancestor (Wolf et al. 2018). Thus, continuing to expand our understanding of the relatedness of xrRNA folds will not only inform computational searches for additional versions, but will open the possibility to use the similarity of structured RNA elements with conserved functional roles as another criterion for virus relatedness, evolution, and taxonomy.

## DATA DEPOSITION

The coordinates have been deposited in the Protein Data Bank with the accession code XXXX (to be supplied prior to publication)

## ACKNOWLEDGMENTS

We wish to thank Pascal Auffinger for helpful suggestions during refinement. Support was provided by the National Institutes of Health (Grants R35GM118070 and R01AI133348 to J.S.K) and the Deutsche Forschungsgemeinschaft Fellowship STE2509/201 to A.-L.S.). Beamline 4.2.2 at the Advanced Light Source is partially funded by the National Institutes of Health (P30GM124169-01) and operated under contract with the U.S. Department of Energy (DE-AC02-05CH11231). The University of Colorado Anschutz Medical Campus x-ray crystallography facility is supported by the National Institutes of Health (Grants P30CA046934 and S10OD012033).

## MATERIALS AND METHODS

### DNA templates and mutagenesis

DNA templates for *in vitro* transcription were gBlocks ordered from IDT, cloned into pUC19 and verified by sequencing. RNA constructs for Xrn1 degradation assays contained the xrRNA sequence plus 34 nucleotides of the endogenous upstream sequence of the viral genome (‘leader sequence’) to allow loading of the exoribonucleases. Nucleotide numbering throughout the manuscript is in reference to the crystallized RNA construct. Transcription with T7 RNA polymerase was started with 2 guanosine nucleotides for enhanced transcription efficiency.

### *In vitro* transcription

DNA templates for *in vitro* transcription were amplified by PCR using custom DNA primers (IDT) and Phusion Hot Start polymerase (New England BioLabs). 2.5 mL transcription reactions were assembled using 1000 µL PCR reactions as template (∼0.2 µM template DNA), 6 mM each NTP, 60 mM MgCl_2_, 30 mM Tris pH 8.0, 10 mM DTT, 0.1% spermidine, 0.1% Triton X-100, T7 RNA polymerase and 2 µL RNasin RNase inhibitor (Promega) and incubated overnight at 37°C. After inorganic pyrophosphates were precipitated by centrifugation, the reactions were ethanol precipitated and purified on a 7 M urea 8% denaturing polyacrylamide gel. RNAs of the correct size were gel-excised, eluted overnight at 4°C into ∼40 mL of diethylpyrocarbonate (DEPC)-treated milli-Q filtered water (Millipore) and concentrated and washed using Amicon Ultra spin concentrators (Millipore).

### Protein expression and purification

The expression vector for *Kluyveromyces lactis* Xrn1 (Chang et al. 2011) (residues 1-1245) was a gift of Prof. Liang Tong at Columbia University and the expression vector for *Bdellovibrio bacteriovorus* RppH was a gift of Joel Belasco at NYU (Messing et al. 2009). All recombinant proteins were 6XHis-tagged, expressed in *E. coli* BL21 cells and purified using Ni-NTA resin (Thermo), followed by size exclusion with either a Superdex 75 or Superdex 200 column in an AKTA pure FPLC (GE Healthcare). The final product was stored in buffer containing 20 mM Tris pH 7.3, 300 mM NaCl, 1 mM DTT or 2 mM BME, and 10% glycerol at −80°C. The purity of the recombinant proteins was verified by SDS-PAGE and Coomassie staining.

### 5’-3’ exoribonuclease degradation assay

4 µg RNA was resuspended in 40 µL 100 mM NaCl, 10 mM MgCl_2_, 50 mM Tris pH 7.5, 1 mM DTT and re-folded at 90°C for 3 minutes then 20°C for 5 minutes. 3 µL recombinant RppH (0.5 µg/µL stock) was added and the samples were split into two 20 µL reactions (-/+ exoribonuclease). 1.5 µL of the recombinant Xrn1 (0.8 µg/µL stock) was added where indicated. All reactions were incubated for 1.5 hrs at 30°C using a thermocycler. The degradation reactions were resolved on a 7 M urea 10% denaturing polyacrylamide gel and stained with ethidium bromide.

### ^32^P-5’-end labeling of RNA

RNA was dephosphorylated using rApid Alkaline phosphatase (Millipore) to convert 5’-triphosphate to 5’-monophosphate ends. To this end, 100 pmol RNA was incubated in a 20 μl reaction containing 2 μl rApid Alkaline phosphatase buffer, 2 μl rApid Alkaline phosphatase, 1 μl RNasin Ribonuclease Inhibitor (Sigma-Aldrich), and incubated for 30 mins at 37°C. The enzyme was heat-inactivated for 2 mins at 75°C. The RNA was subsequently 5’-end labeled with [γ32-P] ATP (PerkinElmer) using T4 polynucleotide kinase (PNK) (New England Biolabs). This was done by adding 1 μl 100mM MgCl_2_, 4 μl PNK Buffer, 2 μl of 5 mCi [γ32-P] ATP, and 2 μl of PNK in a total volume of 40 μl. The reaction mix was incubated for 30 mins at 37°C and purified using Micro-Bio P-30 spin columns (Bio-Rad).

### Electrophoretic mobility shift assay

Unlabeled RNA of the indicated concentrations was diluted in 8 µL 1X TH-loading buffer (66 mM Tris-HCl, 34 mM HEPES, 3 mM MgCl_2,_ 50% glycerol, xylene cyanol, bromophenol blue) supplemented with trace amounts of ^32^P-labeled RNA, heated for 3 min at 85°C, cooled to room temperature and loaded on an 8% 1x TH-polyacrylamide native gel. The gels were run at 70V in the cold room in 1x TH buffer (66 mM Tris-HCl, 34 mM HEPES, 3 mM MgCl_2_) until bromophenol blue front reached the lower quarter of the gel (∼3 hours), dried and imaged using a phosphor screen and Typhoon 9400 scanner (GE Life Sciences).

### RNA crystallization and diffraction data collection

RNA for crystallization was prepared as described above. The sequence used for *in vitro* transcription of the RNA that was used for structure determination was 5’-ggAACTAGCtcAGCATACACGAGTTGCAAGCATgGGAAGTTCAAGCCTCGT<u>GGGCGGCAT</u> <u>GGTCCCAGCCTCCTCGCTGGCGCCGCCTGGGCAACATGCTTCGGCATGGCGAATGGGACC</u>-3’ where the underlined sequence belongs to a hepatitis delta ribozyme that was used to generate homogenous 3’ ends. Lowercase letters represent sequences altered to facilitate transcription (two additional G) and the G.U phasing module. Ribozyme cleavage was induced at the end of the transcription reaction by adding MgCl_2_ (final conc. 120 mM) and incubating for 10 min at 65°C. Ribozyme-cleaved RNA was purified on a 7 M urea 8% polyacrylamide gel as described above. 5 mg/mL RNA was re-folded at 65°C for 3 minutes in a buffer containing 30 mM HEPES pH 7.5, 20 mM MgCl_2_, and 100 mM KCl. Crystal Screens I and II, Natrix I and II, and the Nucleic Acid Mini Screen (all from Hampton Research) were used to perform initial screens at 20°C with sitting-drop vapor diffusion crystallization. Initial hits were optimized using custom screens and the Hampton Research Additive Screen.

The RNAs used for the final structural determination were crystallized in drops of 1 μl RNA solution (5 mg/mL RNA in 30 mM HEPES pH 7.5, 20 mM MgCl_2_, 100 mM KCl) + 1 μl crystallization solution (10% 2-methyl-2,4-pentanediol, 40 mM Sodium Cacodylate (pH 6.0), 12 mM Spermine, 150 mM KCl, 100 mM Guanidinium HCl) over a reservoir of 500 μl of 30% 2-methyl-2,4-pentanediol and 100mM Guanidinium HCl. Crystals were buffer-exchanged into freezing solution (30% 2-methyl-2,4-pentanediol, 40 mM Sodium Cacodylate (pH 6.0), 12mM Spermine, 150 mM KCl, 100 mM Guanidinium HCl), with or without 5 mM iridium(III) hexammine to obtain derived crystals for experimental SAD phasing, and then flash-frozen in liquid nitrogen for x-ray diffraction. Diffraction data were collected at Advanced Light Source Beamline 4.2.2 using ‘shutterless’ collection at the Iridium L-I edge (0.9234 Å) at 100°K. Two native and three derivative 180° datasets were collected with 0.2° oscillation images. Data were indexed, integrated, and scaled using XDS (Kabsch 2010). Three datasets collected on the same crystal were used for SAD phasing.

### Structure determination and refinement

Ten iridium(III) hexammine sites were identified and used in SAD phasing within the AutoSol function of Phenix v. 1.13_2998 on Mac OS (overall FOM = 0.56) (Liebschner et al. 2019). The map was used to manually build an initial model, which was improved through iterative rounds of model building and refinement (simulated annealing, Rigid-body, B-factor refinement) using Coot (Emsley et al. 2010) and Phenix. The final model contains two RNA molecules in the asymmetric unit and all 51 nucleotides are resolved.

Peaks in the anomalous map disappearing around 5 sigma were assigned to cacodylate molecules (f” (As) ∼ 3 electrons at 0.9234 Å), also taking geometry into account. The model was refined in Phenix v. 1.17.1_3660 on Mac OS, using the graphic user interface, down to R/R_free_ of 0.25/0.28. The GTP introduced upon transcription at the 5’ end and the 2’,3’ cyclic UMP introduced during ribozyme cleavage at the 3’ end were linked to the RNA chain using JLigand in CCP4i v. 7.0 (Lebedev et al. 2012; Winn et al. 2011). Map calculation in Refmac v. 5.8.0257 within CCP4i (Winn et al. 2011) led to a higher-quality map with clear density peaks in both 2Fo-Fc and Fo-Fc for the guanidinium ligands. The structure was further refined in Refmac to R/R_free_ 0.23/0.27. The deposited structure (R/R_free_ of 0.229/0.258) was obtained from phenix.refine using default parameters for TLS refinement, occupancy refinement, and optimization of X-ray/stereochemistry and X-ray/B-factor weights. The geometry was validated using the Molprobity web server (Williams et al. 2018).

## SUPPLEMENTAL MATERIALS

**Table 1.** Data collection and refinement statistics.

|  |  |
| --- | --- |
| **Wavelength** | 0.9235 |
| **Resolution range** | 49.72 - 2.60 (2.697 - 2.60) |
| **Space group** | P 4<sub>3</sub> 2<sub>1</sub> 2 |
| **Unit cell** | 72.19 72.19 137.14 90 90 90 |
| **Total reflections** | 482755 (43140) |
| **Unique reflections** | 11689 (1135) |
| **Multiplicity** | 41.3 (38.0) |
| **Completeness (%)** | 99.64 (99.56) |
| **Mean I/sigma(I)** | 30.86 (2.53) |
| **Wilson B-factor** | 63.30 |
| **R-merge** | 0.09956 (1.663) |
| **R-meas** | 0.1008 (1.685) |
| **R-pim** | 0.01561 (0.2697) |
| **CC<sub>1/2</sub>** | 1 (0.914) |
| **CC\*** | 1 (0.977) |
| **Reflections used in refinement** | 11669 (1134) |
| **Reflections used for R-free** | 598 (53) |
| **R-work** | 0.2293 (0.4010) |
| **R-free** | 0.2580 (0.4197) |
| **CC(work)** | 0.945 (0.685) |
| **CC(free)** | 0.910 (0.774) |
| **Number of non-hydrogen atoms** | 2340 |
| **macromolecules** | 2144 |
| **ligands** | 196 |
| **RMS(bonds)** | 0.002 |
| **RMS(angles)** | 0.54 |
| **Rotamer outliers (%)** | 0.00 |
| **Clashscore** | 4.07 |
| **Average B-factor** | 64.64 |
| **macromolecules** | 59.80 |
| **ligands** | 117.62 |
| **Number of TLS groups** | 1 |

Statistics for the highest-resolution shell are shown in parentheses.

**Supplemental Fig. S1:**
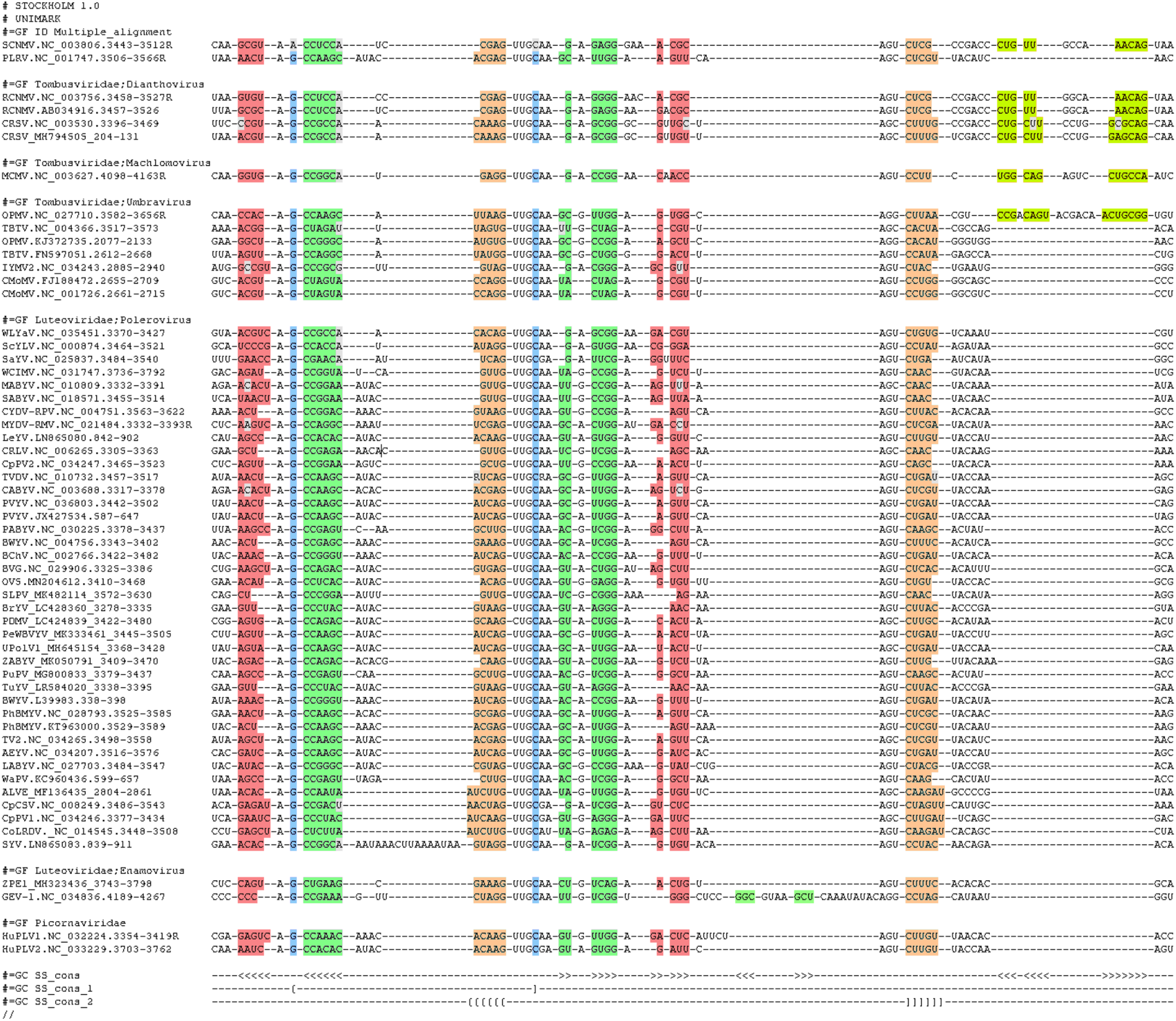
Alignment of xrRNA_LT_ sequences. Each entry lists the official virus acronym, the NCBI accession code, the location of the xrRNA and the sequence in Stockholm format. The alignment in (Steckelberg and Vicens et al. 2018b) was expanded with viral sequences added to the NCBI database between December 2018 and December 2019. The .sto alignment file is also available as a supplemental file. P1, P2, P3, P4 shown in red, green, orange, and lime, respectively. The G–C pair present in Polerovirus but not SCNMV is highlighted in blue.

**Supplemental Fig. S2:**
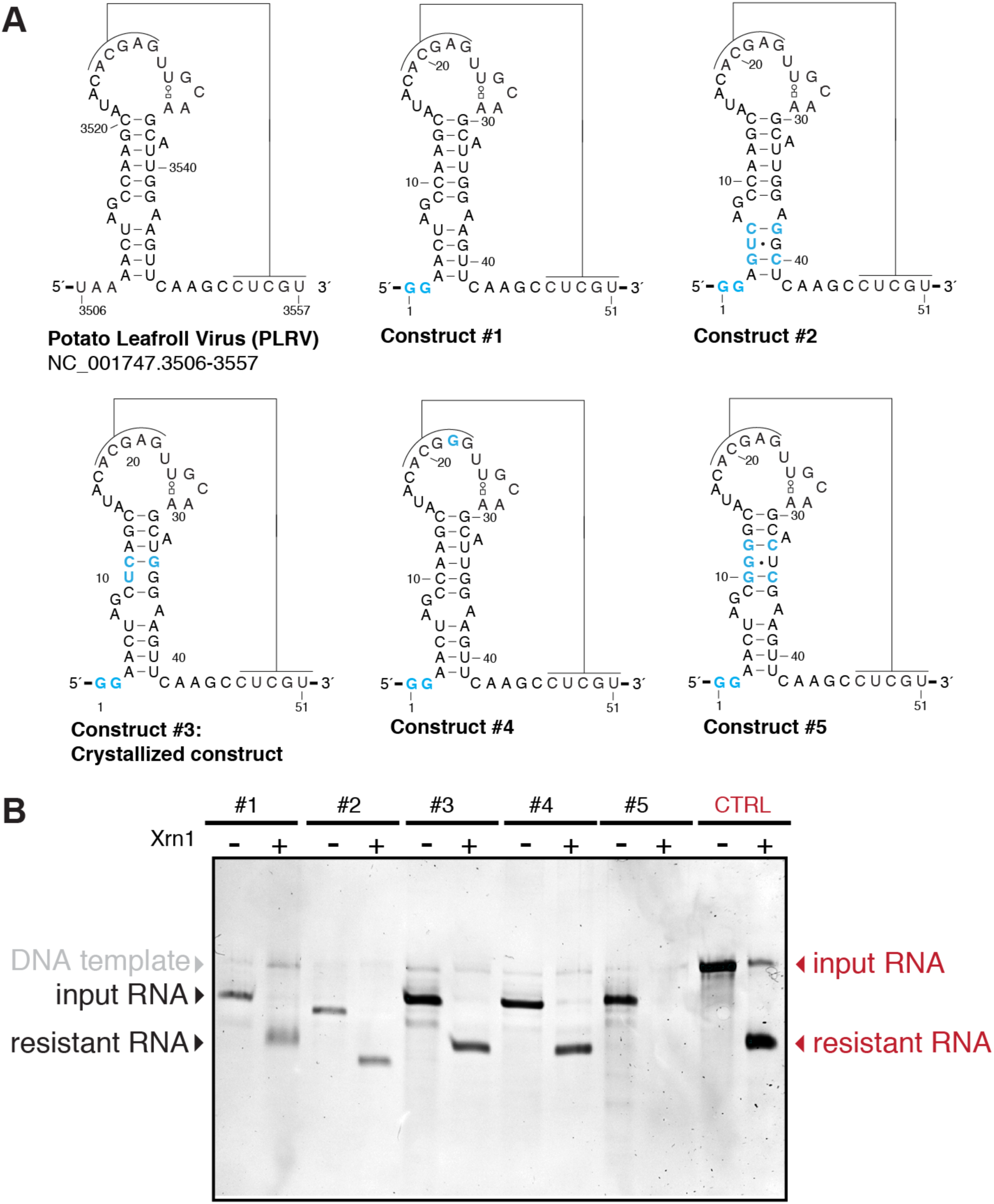
PLRV crystal constructs used in this study. (A) Wild-type sequence for the PLRV xrRNA_LT_ and the five crystal constructs tested for this study. Constructs #2–5 incorporate the G.U phasing module in various locations (Keel et al. 2007). (B) Xrn1 resistance assays of constructs #1–5. Digestion occurred for 2 h at 30°C, as described (Steckelberg and Vicens et al. 2018b). Denaturing PAGE stained with ethidium bromide. The leader was 15 nucleotides long for #1–5 and 35 nucleotides long for the control sequence (wild-type PLRV).

**Supplemental Fig. S3:**
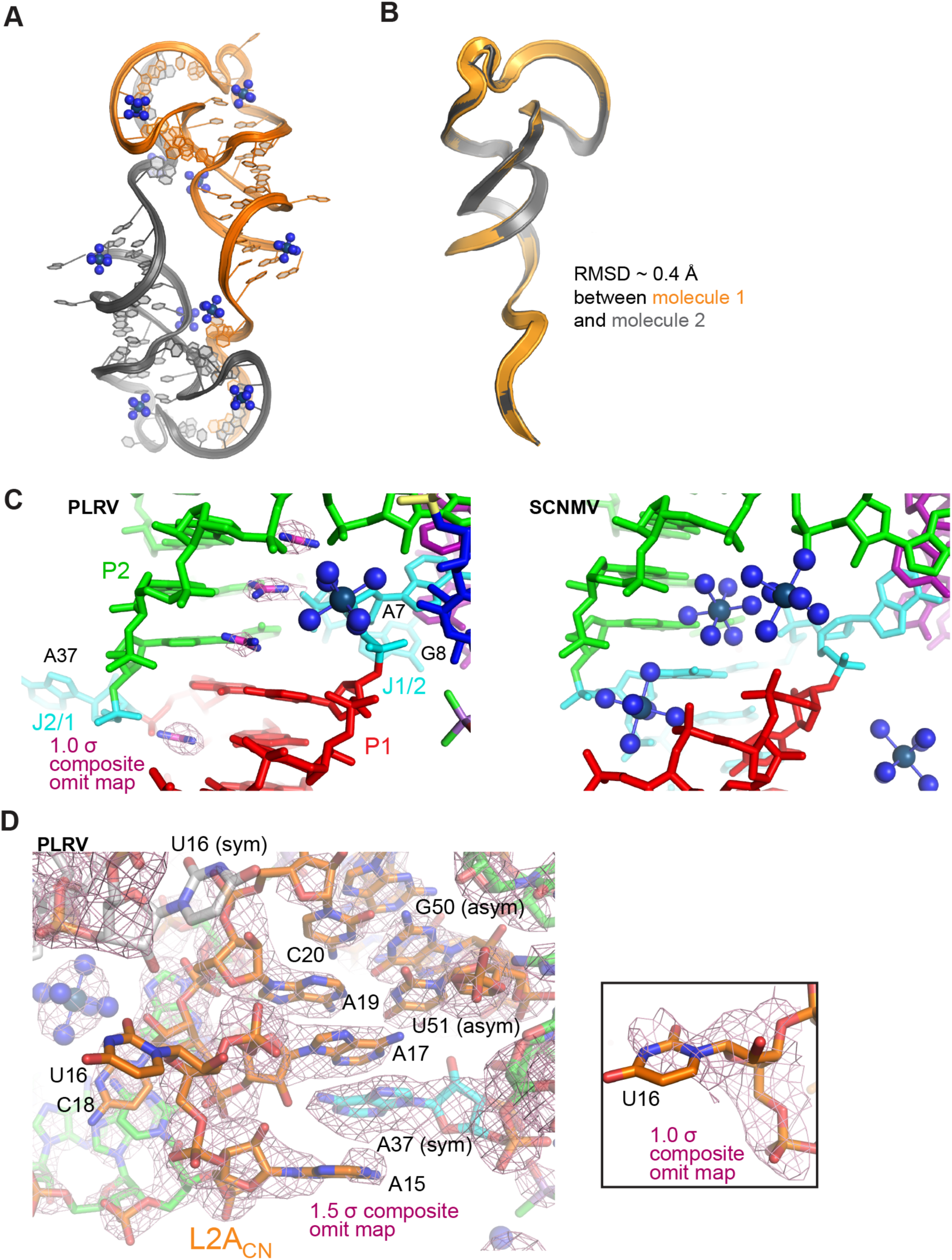
Structural features of the PLRV xrRNA_LT_. (A) Two xrRNAs form a dimer in the asymmetric unit. Blue, iridium(III) hexammine molecules used for experimental phasing. (B) Structural alignment of molecules 1 and 2 in the asymmetric unit (RMSD ∼ 0.4 Å). (C) Guanidinium molecules (carbon atom in magenta) bind near the J1/2 region of PLRV (left), where iridium(III) hexammine ligands bind in the SCNMV structure (right). (D) Stacking, base-pairing and other structural features at the L2A_CN_ – J1/2 – J1/3_PK_ interface. Nucleotides belonging to the second molecule in the asymmetric unit or to symmetry-related molecules in the crystals are denoted by asym and sym, respectively. The composite 2Fo-Fc map shown in panels (C) and (D) was calculated in Phenix using default parameters.

**Supplemental Fig. S4:**
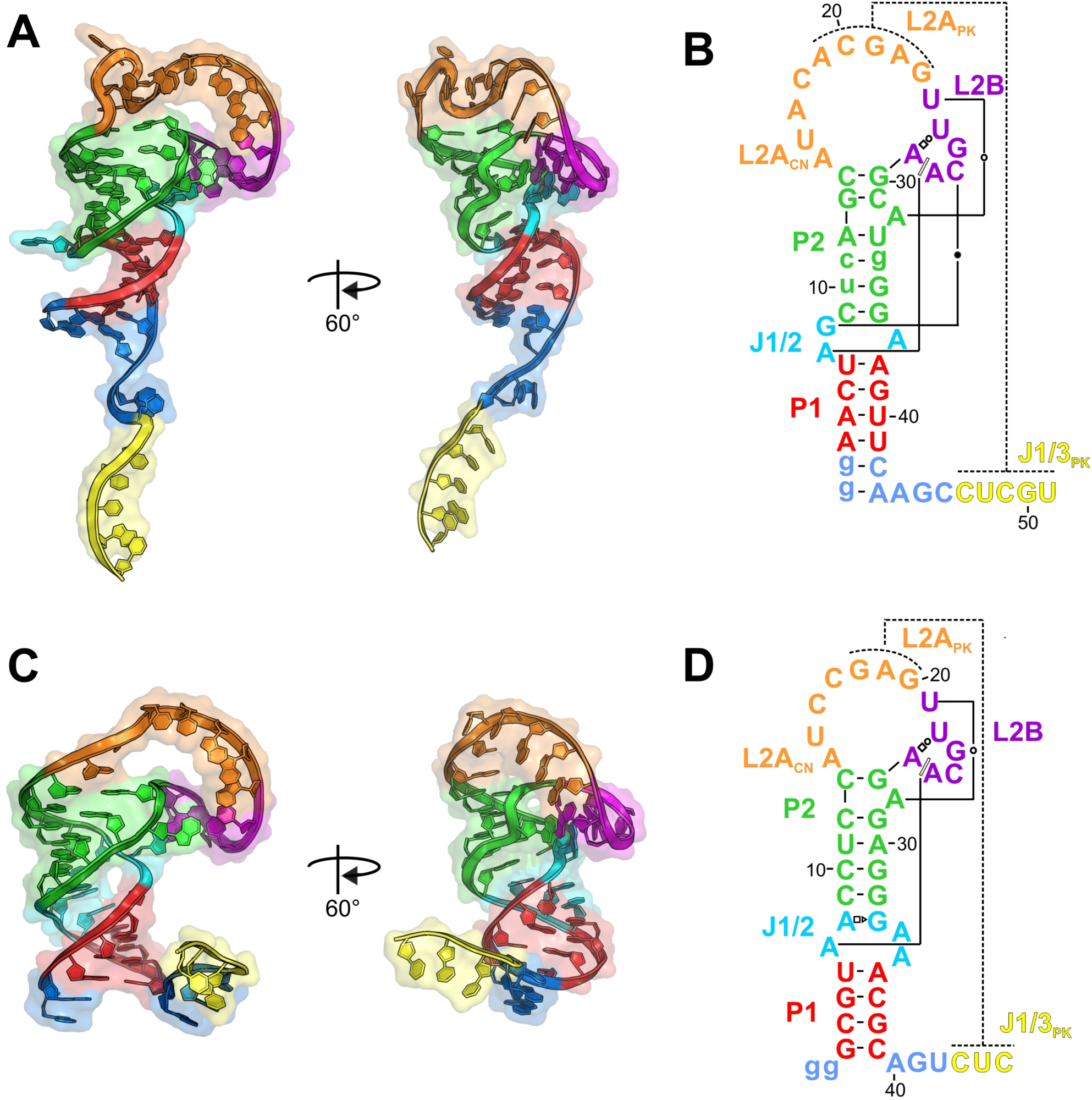
Comparison of PLRV and SCNMV xrRNA structures. (A) Ribbon representation of the PLRV xrRNA structure. (B) Secondary structure of the crystallized PLRV RNA. (C) Ribbon representation of the SCNMV xrRNA. (D) Secondary structure of the SCNMV RNA. Colors match Fig 2. Engineered nucleotides are shown in lowercase letters. Non–Watson–Crick base pairs are in Leontis–Westhof annotation (Leontis and Westhof 2001).

**Supplemental Fig. S5:**
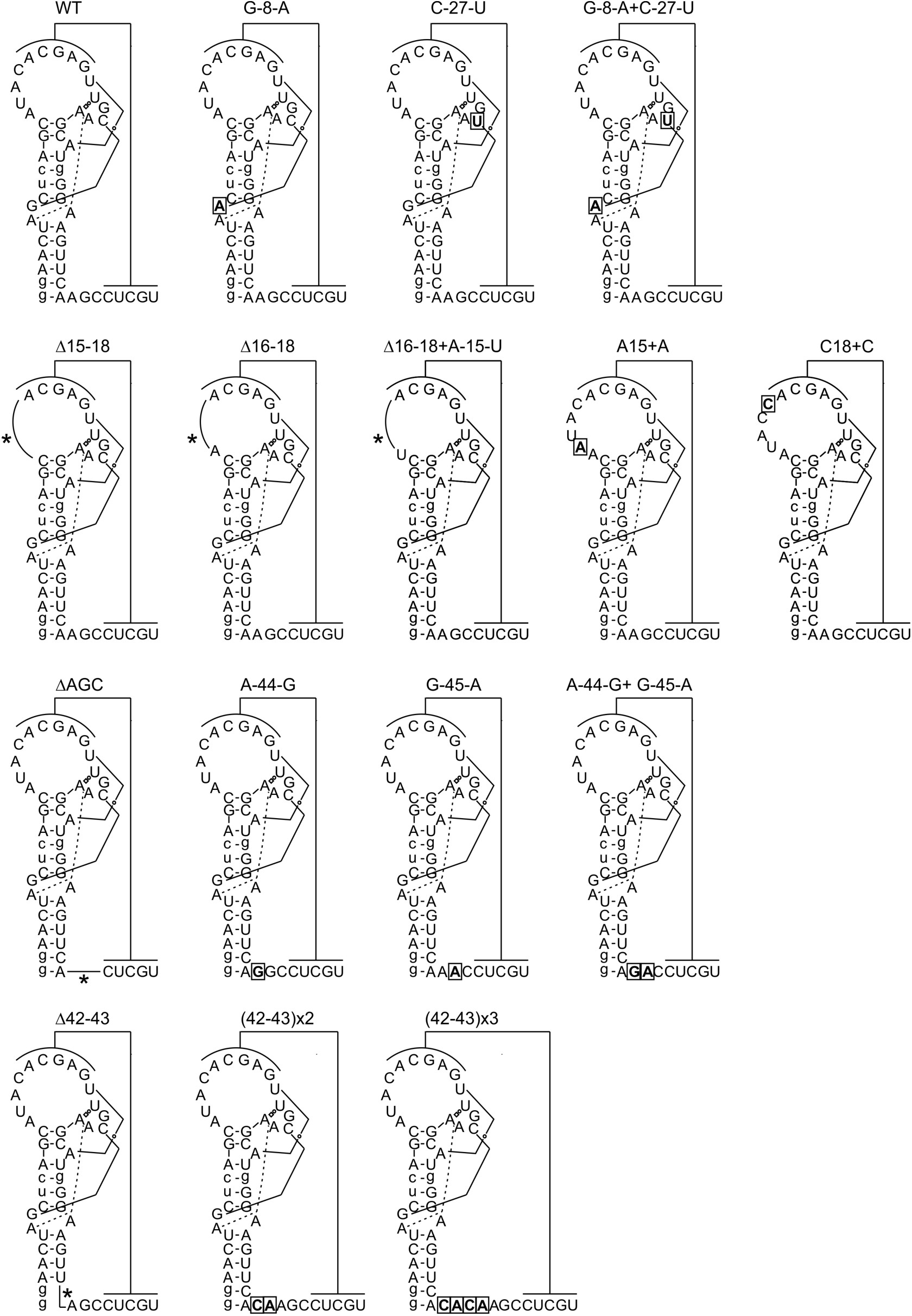
Predicted secondary structure diagrams for the mutants used in functional assays. Secondary structures of wild-type and mutant PLRV xrRNA constructs used in *in vitro* assays. Substitution and insertion mutations are shown boxed and in bold, deletion mutations are shown as lines and indicated by an asterisk (*). For all RNAs, the sequence shown was preceded by a 34 nucleotide leader based on the endogenous viral genomic context (5’-GCCACCACAAAAGAACACUG AAGGAGCUCACUAA-3’), to allow efficient loading of Xrn1.

**Supplemental Fig. S6:**
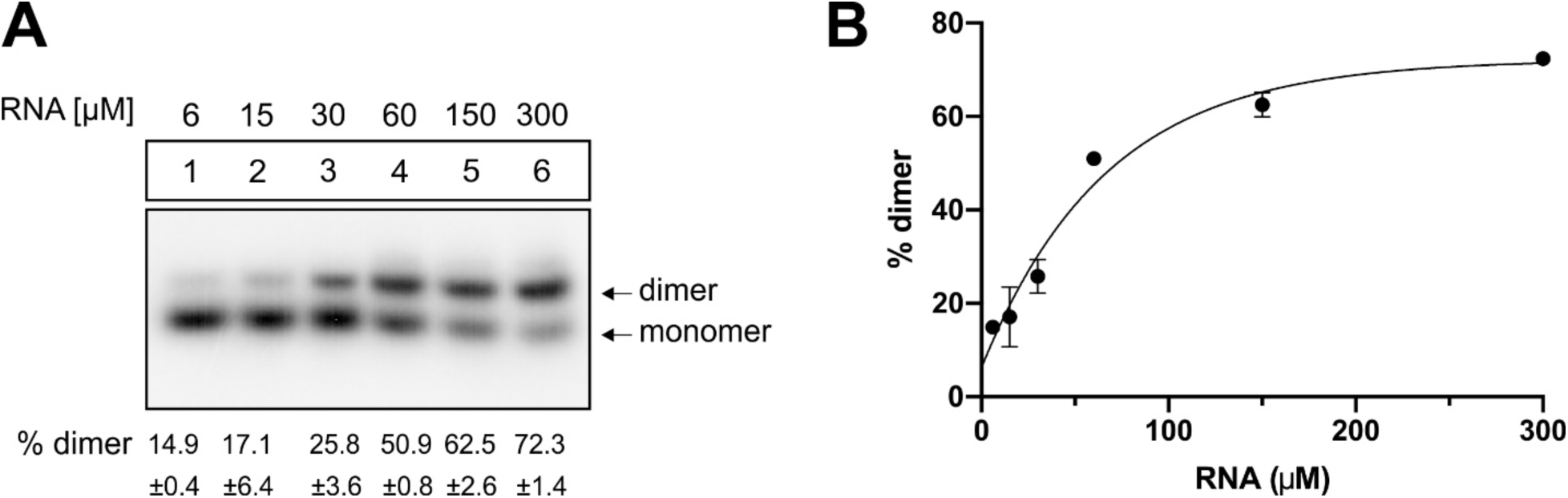
The PLRV xrRNA dimerizes at high concentration. (A) Electron mobility shift assay (EMSA) of the PLRV RNA used for crystallization and structure determination. Numbers under the gel represent the average of 3 independent experiments −/+ standard deviation. (B) Single-exponential fitting of EMSA data from (A). 50% of the RNA is dimerized at 46 μM.

